# A comprehensive proteome and the first phosphoproteome reveal extensive phosphorylation of carbohydrate metabolism in *Cryptosporidium parvum* sporozoites

**DOI:** 10.64898/2026.01.20.700507

**Authors:** Dongqiang Wang, Meng Li, Chenchen Wang, Haitao Li, Jigang Yin, Guan Zhu

## Abstract

*Cryptosporidium parvum* is an obligate intracellular apicomplexan parasite and a major cause of diarrheal disease in humans and animals worldwide. Despite its public health importance, the molecular regulation of parasite metabolism, particularly at the post-translational level, remains poorly understood. Here, we present a comprehensive proteomic analysis and the first phosphoproteomic profile of excysted *C. parvum* sporozoites, the invasive stage responsible for host cell entry. Using data-independent acquisition–based mass spectrometry, we identified 2,272 proteins, representing approximately 58% of the predicted parasite proteome, and 8,994 phosphorylation sites across 833 phosphoproteins. Comparative analyses revealed weak correlations between transcript and protein abundance, underscoring extensive post-transcriptional regulation in sporozoites. Functional enrichment analyses showed that proteins involved in carbohydrate metabolism, particularly glycolysis, are highly abundant in the sporozoite proteome. In contrast, phosphoproteomic data revealed that many core glycolytic enzymes exhibit relatively low phosphorylation propensity, suggesting limited reliance on phosphorylation-based regulation for basal energy metabolism at this stage. To integrate proteomic and phosphoproteomic measurements acquired independently, we developed a relative phosphorylation index (RPI) that enables comparative assessment of phosphorylation propensity across proteins. Application of this metric highlighted selective phosphorylation of proteins associated with signaling, cytoskeletal organization, and host–parasite interaction, while key metabolic entry-point enzymes, such as hexokinase, showed high abundance but minimal detectable phosphorylation, in contrast to their mammalian counterparts. Together, these findings provide the most comprehensive molecular resource to date for *C. parvum* sporozoites and reveal a phosphorylation landscape that emphasizes regulatory and structural processes over core metabolic flux. This work establishes a foundation for understanding parasite-specific regulatory strategies and may inform the identification of novel therapeutic targets against cryptosporidiosis.

**Author Summary:** *Cryptosporidium parvum* is a microscopic parasite that causes diarrheal disease in humans and animals worldwide, yet we still know surprisingly little about how it regulates its biology at the molecular level. In this study, we set out to better understand how the parasite prepares for infection by examining the proteins present in sporozoites, the life stage that actively invades host cells. We generated the most comprehensive protein catalog to date for *C. parvum* sporozoites and, for the first time, mapped thousands of chemical modifications known as protein phosphorylation. These modifications often act as molecular switches that control how proteins behave. We found that proteins involved in basic energy production are extremely abundant but show relatively little phosphorylation, while proteins linked to cell structure, movement, and interaction with the host are more frequently modified. This suggests that the parasite prioritizes regulation of invasion-related processes over fine-tuning its core metabolism at this stage. By providing a detailed molecular resource and a new way to compare protein phosphorylation patterns, our work lays the groundwork for future studies of parasite biology and may help guide the search for new strategies to combat cryptosporidiosis.

## Introduction

*Cryptosporidium parvum* is a unicellular enteric parasite in the phylum Apicomplexa of major medical and veterinary importance [1,2]. It is a frequent cause of waterborne and foodborne outbreaks of cryptosporidiosis worldwide [3,4]. Individuals with immature or compromised immune systems, including infants, young children, and the elderly, are particularly vulnerable to infection [5]. In immunocompromised patients, such as those with AIDS, cryptosporidiosis can become chronic and life-threatening [6]. In livestock, *C. parvum* is a common cause of neonatal diarrhea in calves, lambs, and goat kids, leading to substantial economic losses in the animal production industry [7,8]. The parasite is transmitted via the fecal–oral route, with environmentally resistant oocysts serving as the infectious stage [1]. Following ingestion by a new host, oocysts undergo excystation in response to elevated temperature and bile salts in the small intestine, releasing sporozoites. These excysted sporozoites penetrate the mucosal layer and invade intestinal epithelial cells, initiating a complex intracellular life cycle that includes both asexual and sexual development and culminates in the production of new oocysts.

Despite its importance to human and animal health and substantial progress over the past decade, our understanding of *Cryptosporidium* biology remains limited, largely due to difficulties in obtaining sufficient parasite material and in experimentally manipulating the parasite both in vitro and in vivo [1,9,10]. In the context of omics research, most studies have focused on genomics and transcriptomics, for which DNA and RNA can be amplified from limited starting material [11–18]. In contrast, proteomic analyses require comparatively large amounts of purified parasite material and are therefore less common. To date, only a limited number of sizeable proteomic studies have been reported for *C. parvum* sporozoites or oocysts, stages for which relatively pure parasite material can be obtained in quantity [19–25]. These studies identified between 135 or 303 and 1,237 proteins [21–23]. A more recent hyperLOPIT-based spatial proteomic analysis assigned 1,107 sporozoite proteins to 14 subcellular compartments [20], while an enriched oocyst wall proteome identified 798 proteins [24]. In addition, a proteomic study of *Cryptosporidium andersoni*, a bovine gastric species, identified 1,586 proteins [25]. Given that the *C. parvum* genome is predicted to encode 3,944 proteins based on the original annotation of the Iowa II strain [26], or 3,896 proteins based on reannotation of the Iowa-ATCC strain [27], existing proteomes collectively represent no more than approximately one-third of the predicted proteome.

Notably, global post-translational modification (PTM) proteomes have not been reported for *Cryptosporidium*. Previous PTM-related studies have focused primarily on glycoproteins, using chemical deglycosylation, lectin binding, or glycan-specific antibodies to validate glycosylation of selected native proteins such as Gp900 and Gp40 [28,29]. Three small-scale mass spectrometry–based glycoproteomic studies further characterized glycan profiles of a limited number of N- and O-linked glycoproteins [30,31] and identified N-, O-, and C-glycosylation sites on seven thrombospondin repeat–containing proteins [32]. Beyond glycosylation, however, other PTMs—including phosphorylation, ubiquitination, acetylation, and methylation—remain essentially unexplored in *C. parvum*. These modifications are known to substantially expand protein functional diversity by modulating enzymatic activity, stability, localization, and protein–protein interactions [33–35]. The absence of PTM-scale datasets therefore represents a major gap in our understanding of post-translational regulation in *Cryptosporidium*.

Here, we report an integrated mass spectrometry–based proteomic and phosphoproteomic analysis of excysted *C. parvum* sporozoites, the invasive stage of the parasite. The proteome identifies 25,506 peptides mapped to 2,272 proteins, representing approximately 58% of the predicted *C. parvum* proteome. In parallel, phosphoproteomic profiling identifies 5,690 phosphopeptides mapped to 2,141 proteins, providing the first global view of protein phosphorylation in *C. parvum*. Phosphorylation predominantly occurs on serine residues, followed by threonine and tyrosine, with distinct sequence motifs and subcellular distribution patterns supported by immunofluorescence analysis using pan-phosphoprotein antibodies. To facilitate comparative analysis between independently acquired proteome and phosphoproteome datasets, we introduce a relative phosphorylation index to assess phosphorylation propensity relative to protein abundance. Integrative pathway analyses reveal that carbohydrate metabolism, including glycolysis and connected pathways, is highly represented in both the proteome and phosphoproteome, with multiple enzymes exhibiting disproportionately high phosphorylation propensity. Together, these data establish the largest sporozoite proteome and the first phosphoproteome of *C. parvum*, providing a systems-level framework for understanding protein expression and phosphorylation associated with metabolic activity in the invasive stage of the parasite.

## Materials and Methods

### Ethics statement

Animal experiments were conducted in compliance with the Guide for the Care and Use of Laboratory Animals of the Ministry of Health of China and were approved by the Animal Welfare and Research Ethics Committee of the Institute of Zoonosis, Jilin University (approval no. #2024-IZ-00039).

### Parasite materials and sporozoite preparation

The JLU01 isolate of *Cryptosporidium parvum* (gp60 subtype IIdA19G1) was propagated in bovine calves as described previously [6,36]. Oocysts of *C. parvum* were purified from calf feces by discontinuous sucrose/CsCl gradient centrifugation and stored in phosphate-buffered saline (PBS) containing penicillin (200 U/mL) and streptomycin (0.2 mg/mL) at 4 °C [6,37,38].

This study focuses on the proteome and phosphoproteome of *C. parvum* sporozoites, the invasive stage of the parasite, for which relatively pure parasite material can be obtained in sufficient quantity. Free sporozoites were prepared using an in vitro excystation protocol, with oocysts stored for less than two months after harvest. Prior to excystation, oocysts were treated with 10% household bleach for 5 min on ice to inactivate residual bacterial contaminants, followed by 5–8 washes with PBS.

Excystation was performed by incubating oocysts in RPMI-1640 medium supplemented with 0.5% taurocholic acid sodium salt at 37 °C for 45 min, yielding an excystation efficiency of >90%. Excysted sporozoites were washed three times with RPMI-1640 medium prewarmed to 37 °C to minimize heat or cold shock, using centrifugation at 10,000 × g for 5 min. After each centrifugation step, microtubes were gently tapped to loosen the pellets prior to resuspension in the medium.

After the final wash, sporozoites (4 × 10^8^ per sample replicate) were resuspended in Pierce RIPA buffer (Thermo Fisher Scientific) containing a protease inhibitor cocktail for eukaryotes (Sigma-Aldrich) and lysed by three cycles of freeze–thawing in liquid nitrogen and on ice. Lysates were centrifuged at 10,000 × g for 10 min at 4 °C, and the supernatants were collected, snap-frozen in liquid nitrogen, and shipped on dry ice to BGI Genomics (Shenzhen, Guangdong, China) for proteomic analysis and to PTM Bio (Hangzhou, Zhejiang, China) for phosphoproteomic analysis, as described below.

### LC–MS/MS-based proteome analysis

#### High-pH reversed-phase fractionation

At BGI Genomics, the quantity and quality of protein extracts were evaluated by Bradford assay and SDS–PAGE, followed by digestion with trypsin in 50 mM NH₄HCO₃ (enzyme/protein ratio = 1:40, w/w) at 37 °C for 4 h. Digested samples were desalted using a Strata X column (Phenomenex, Torrance, CA, USA) and vacuum-dried prior to high-pH reversed-phase fractionation by high-performance liquid chromatography (HPLC). Samples were dissolved in mobile phase A (5% acetonitrile, pH 9.8) and separated on a Gemini C18 high-pH column (5 μm, 4.6 × 250 mm; Phenomenex) using a Shimadzu LC-20AB HPLC system. Peptides were eluted at a flow rate of 1 mL/min with a gradient of mobile phase B (95% acetonitrile, pH 9.8) as follows: 5% B for 10 min; 5–35% B over 40 min; 35–95% B over 1 min; and 95% B for 3 min, followed by re-equilibration at 5% B for 10 min. Elution was monitored at 214 nm, and fractions were collected at 1-min intervals. Collected fractions were subsequently combined into 10 pooled samples and dried by vacuum centrifugation for downstream analysis.

#### DDA and DIA analysis by nano-LC–MS/MS

Dried peptide samples were reconstituted in mobile phase A (2% acetonitrile, 0.1% formic acid), centrifuged at 20,000 × g for 10 min, and the clarified supernatants were used for analysis. Peptide separation was performed on a Thermo UltiMate 3000 UHPLC system. Samples were first loaded onto a trap column for enrichment and desalting and then separated on a self-packed C18 analytical column (150 μm inner diameter, 1.8 μm particle size, 35 cm length). Peptides were eluted at a flow rate of 500 nL/min using the following gradient of mobile phase B (98% acetonitrile, 0.1% formic acid): 0–5 min, 5% B; 5–130 min, 5–25% B; 130–150 min, 25–35% B; 150–160 min, 35–80% B; 160–175 min, 80% B; 175–175.5 min, 80–5% B; and 175.5–180 min, 5% B. The nano-LC eluent was directly introduced into a Fusion Lumos mass spectrometer (Thermo Fisher Scientific) via nano-electrospray ionization.

For data-dependent acquisition (DDA), peptides were analyzed in positive-ion mode with a spray voltage of 2 kV. MS¹ spectra were acquired in the Orbitrap at a resolution of 60,000 over an m/z range of 350–1,500, with a maximum injection time (MIT) of 50 ms and an automatic gain control (AGC) target of 3 × 10^6^. The top 30 precursor ions (intensity >2 × 10^4^; charge states 2+ to 6+) were selected for higher-energy collisional dissociation (HCD; normalized collision energy [NCE] 30). MS/MS spectra were acquired at a resolution of 15,000 with an MIT of 50 ms and an AGC target of 1 × 10^5^. Dynamic exclusion was set to 30 s, and MS/MS scans were acquired starting at m/z 100.

For data-independent acquisition (DIA), peptides were analyzed under the same nano-electrospray conditions using a spray voltage of 2 kV. MS¹ spectra were acquired at a resolution of 60,000 over an m/z range of 400–1,500 with an MIT of 50 ms. The m/z range was partitioned into 44 consecutive DIA windows for MS/MS acquisition. Fragmentation was performed by HCD (NCE 30), and MS/MS spectra were acquired in the Orbitrap at a resolution of 30,000 with an MIT of 54 ms and an AGC target of 5 × 10^4^.

DDA data were identified using the Andromeda search engine implemented in MaxQuant, and the identification results were used for spectral library construction. For large-scale DIA data, the mProphet algorithm was applied for analytical quality control, yielding a large number of reliable quantitative measurements.

#### Proteomics data processing and database searching

Peptide and protein identification were performed using a combined DDA–DIA workflow following the facility’s standard bioinformatics pipeline. For spectral library generation, DDA raw files were processed with MaxQuant (Max Planck Institute) using the Andromeda search engine. MS/MS spectra were searched against the NCBI RefSeq protein database for *C. parvum*, supplemented with a reverse decoy database for false discovery rate (FDR) estimation. Trypsin/P was specified as the digestion enzyme, with up to two missed cleavages permitted. Carbamidomethylation of cysteine was set as a fixed modification, and oxidation of methionine and N-terminal acetylation were included as variable modifications. Precursor and fragment mass tolerances were set according to instrument specifications, and peptide-spectrum matches (PSMs), peptides, and proteins were filtered to a 1% FDR. The resulting high-confidence identifications were used to construct a project-specific spectral library.

DIA raw files were analyzed using the facility’s integrated DIA workflow, which applies indexed retention-time (iRT) calibration followed by targeted data extraction using the spectral library. Peak detection, scoring, and FDR control were performed using an mProphet-based statistical model, with peptide- and protein-level FDR thresholds set at 1%. For samples that passed quality-control criteria, peptide intensities were integrated and normalized to generate protein-level quantitative matrices. The final output included peptide- and protein-level abundance tables (designated “DIA-peptides” and “DIA-proteins,” respectively), each containing RefSeq accession numbers and corresponding quantitative values.

RefSeq accession numbers in the final datasets were further mapped to gene identifiers assigned to the *C. parvum* Iowa II reference genome available in CryptoDB (https://cryptodb.org).

### LC–MS/MS-based phosphoproteome analysis

#### Sample preparation and phosphopeptide enrichment

Free sporozoites were prepared by excystation as described above. To maximize detection sensitivity using the available parasite material, three independently prepared sporozoite samples (4 × 10^8^ sporozoites per sample) were pooled for phosphoproteomic analysis. Proteins were precipitated by adding one volume of pre-chilled acetone with vortexing, followed by four additional volumes of cold acetone, and incubated at −20 °C for 2 h. Protein precipitates were washed 2–3 times with cold acetone, briefly air-dried, and resuspended in 200 mM triethylammonium bicarbonate (TEAB) by ultrasonic dispersion.

Proteins were reduced with 5 mM dithiothreitol at 56 °C for 30 min and alkylated with 11 mM iodoacetamide at room temperature for 15 min in the dark. Trypsin was added at an enzyme-to-protein ratio of 1:50 for overnight digestion, and the resulting peptides were desalted using Strata-X solid-phase extraction cartridges.

For phosphopeptide enrichment, peptide mixtures were incubated with immobilized metal affinity chromatography (IMAC) microspheres in loading buffer (50% acetonitrile, 0.5% acetic acid) with gentle agitation. The beads were washed sequentially with 50% acetonitrile/0.5% acetic acid and 30% acetonitrile/0.1% trifluoroacetic acid to remove nonspecifically bound peptides. Phosphopeptides were eluted with 10% ammonium hydroxide, collected, and dried by vacuum centrifugation prior to LC–MS/MS analysis.

#### LC–MS/MS analysis

Dried phosphopeptides were reconstituted in solvent A (0.1% formic acid, 2% acetonitrile in water) and loaded onto a home-packed reversed-phase analytical column (25 cm length, 100 μm inner diameter). Chromatographic separation was performed on a NanoElute UHPLC system (Bruker Daltonics) at a constant flow rate of 450 nL/min using the following gradient of solvent B (0.1% formic acid in acetonitrile): 0–16 min, 2–22% B; 16–22 min, 22–35% B; 22–26 min, 35–90% B; and 26–30 min, 90% B.

Eluted peptides were analyzed on a timsTOF Pro 2 mass spectrometer (Bruker Daltonics) operated in dia-PASEF mode. Ionization was achieved using a capillary electrospray voltage of 1.7 kV. Precursor and fragment ions were analyzed in the time-of-flight (TOF) analyzer. Full MS scans were acquired over an m/z range of 100–1700, and 22 dia-PASEF MS/MS scans were collected per acquisition cycle. MS/MS scans covered an m/z range of 395–1395 using isolation windows of 20 m/z.

#### Database searching and phosphopeptide identification

Raw DIA-PASEF data were processed using Spectronaut (version 18). Spectra were searched against a *C. parvum* RefSeq-based FASTA database concatenated with a reverse decoy database for false discovery rate (FDR) estimation. Trypsin/P was specified as the digestion enzyme, allowing up to two missed cleavages. Carbamidomethylation of cysteine was set as a fixed modification, while phosphorylation of serine, threonine, and tyrosine residues and other relevant post-translational modifications were included as variable modifications. Precursor and fragment mass tolerances were applied according to instrument-specific settings (20 ppm for precursor tolerance in the first and main searches and 0.05 Da for fragment ions). Peptide- and protein-level FDRs were controlled at <1% using Spectronaut’s integrated target–decoy strategy and mProphet-based scoring workflow.

### Bioinformatic analysis

#### Post-translational modification motif analysis

Motifs flanking serine and threonine phosphorylation sites were analyzed using the MoMo (Modification Motifs) tool based on the motif-x algorithm (https://meme-suite.org/meme/tools/momo). The analysis window included six amino acids upstream and downstream of each modification site. A sequence pattern was considered a motif when the number of matching peptides exceeded 20 and the statistical P value was less than 1 × 10^-6^.

In addition to motif logos generated by motif-x, motif heatmaps were produced using a degree of change score (DS), calculated as DS = −log_10_(P value) × sign(diff.percent). P values were derived from Fisher’s exact tests for each amino acid at every position within the MoMo output. The diff.percent term represents the change in frequency of a given amino acid at a specific position relative to the background proteome.

#### Correlation of the sporozoite proteome with existing transcriptomes and proteomes

Previously reported transcriptomic and proteomic datasets were retrieved from CryptoDB, including (i) sporozoite transcriptomes (groups 0 h PBS exp 15 and exp 16) determined by RNA-seq (deposited by Widmer et al.; dataset version 2018-05-22) and (ii) sporozoite proteomes determined by two-dimensional gel LC–MS/MS analysis (deposited by Wastling et al.; dataset version 2007-03-09) [21]. Transcriptomic and proteomic abundance values were log₂-transformed and median-normalized prior to calculation of Spearman correlation coefficients. Spearman correlation analysis was also performed between the sporozoite proteome and phosphoproteome generated in this study.

#### Pathway mapping and comparison of proteome and phosphoproteome

Protein sequences identified in sporozoite proteomic and phosphoproteomic analyses were mapped to Kyoto Encyclopedia of Genes and Genomes (KEGG) pathways using the BlastKOALA tool (https://www.kegg.jp/blastkoala). Ortholog numbers in enriched pathways were compared between proteome and phosphoproteome datasets.

#### Relative phosphorylation index derived from proteome and phosphoproteome datasets

The sporozoite proteome and phosphoproteome of *C. parvum* were acquired and quantified independently, and MS signal intensities in each dataset therefore represent relative abundances that are not directly comparable on an absolute scale. Because experimentally determined phosphorylation stoichiometry (i.e., the fraction of protein molecules that are phosphorylated) is currently unavailable for *C. parvum* proteins and cannot be used for cross-dataset calibration, a comparative metric termed the relative phosphorylation index (RPI) was derived to assess the relative phosphorylation propensity of individual proteins. This approach is conceptually similar to methods that normalize phosphoproteomic measurements by total protein abundance in studies of signaling and deletion strain comparisons to disentangle phosphorylation changes from overall protein expression changes [39,40]. For each dataset, protein abundances (log2-transformed signal intensities) were median-centered by subtracting the dataset-specific median intensity. For proteins quantified in both datasets, RPI was calculated as the difference between median-centered phosphoproteome and proteome intensities, which is equivalent to a phosphoproteome-to-proteome ratio on the linear scale. RPI thus provides a dataset-internal comparative measure that facilitates relative comparisons across proteins while not representing absolute phosphorylation stoichiometry.

### Immunofluorescence assay of phosphorylated proteins in C. parvum sporozoites. **Excysted**

*C. parvum* sporozoites were prepared as described above, washed three times with PBS by centrifugation (10,000 × g for 5 min each), fixed with 4% paraformaldehyde for 30 min, and washed again with PBS. Sporozoites in suspension were placed onto poly-L-lysine–coated glass coverslips, semi-dried, and blocked with 5% bovine serum albumin (BSA).

Sporozoites were incubated with pan-phosphorylated protein antibodies in PBS containing 1% BSA for 1 h. Primary antibodies included anti–phospho-serine rabbit polyclonal antibody (Cat. no. bs-11993R; Bioss, Beijing, China), anti–phospho-threonine rabbit monoclonal antibody (Cat. no. PTM0705RM; PTM Bio, Hangzhou, China), and anti–phospho-tyrosine rabbit monoclonal antibody (Cat. no. PTM0702RM; PTM Bio). After three washes with PBS/1% BSA, samples were incubated with Alexa Fluor 488–conjugated goat anti-rabbit IgG (Thermo Fisher Scientific, West Palm Beach, FL, USA). Samples were washed, counterstained with DAPI (1.0 μg/mL), mounted with antifade mounting medium, and examined using an Olympus BX53 fluorescence microscope. Images were captured with an Olympus DP74 camera and processed uniformly using Adobe Photoshop.

## Results

### Excysted *Cryptosporidium parvum* sporozoites express over half of the predicted proteome

LC–MS/MS–based proteomic analysis of excysted *C. parvum* sporozoites identified a total of 25,506 peptides, including 18,812 unique peptides with lengths ranging from 7 to 47 amino acids (mean = 12.21; peak = 11) (Fig 1A). These peptides mapped to 2,272 proteins, with most proteins exhibiting 10–20% sequence coverage (Fig 1B; Table 1 and Table S1). Protein signal intensities spanned a wide dynamic range, from 10.43 to 25.59 (log2-transformed values), with a median of 16.61 (Fig 1C).

**Figure 1.**
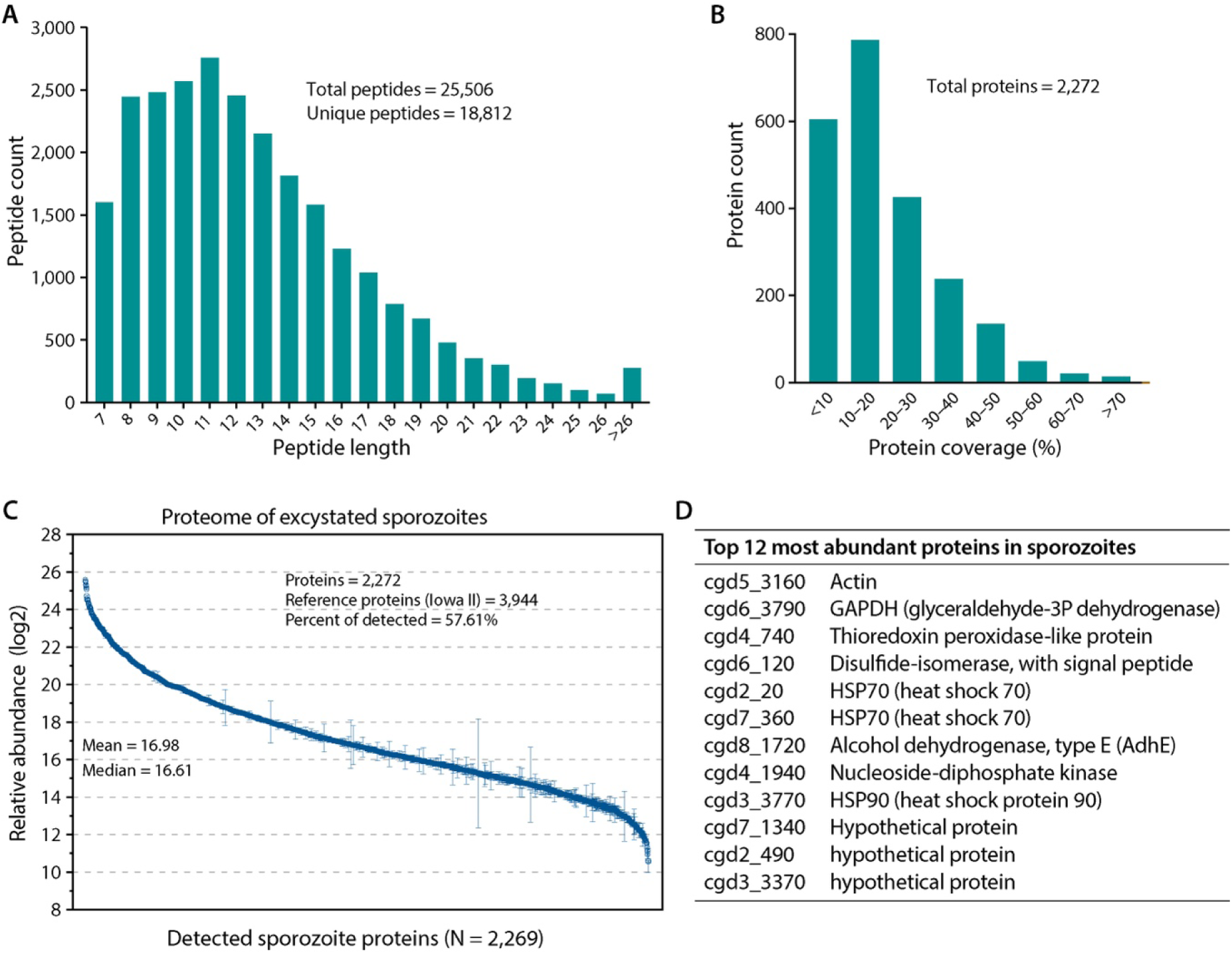
Proteomic profiling of excysted *C. parvum* sporozoites. (A) Distribution of peptide lengths identified in the sporozoite proteome, showing the number of peptides across length intervals. A total of 25,506 peptides, including 18,812 unique peptides, were identified. (B) Protein sequence coverage of identified sporozoite proteins, expressed as the percentage of amino acid residues covered by detected peptides. (C) Distribution of protein abundance values in the sporozoite proteome, shown as log2-transformed intensities. Median and mean values are indicated. (D) The 12 most abundant proteins detected in excysted sporozoites, ranked by proteomic signal intensity. Protein identifiers and annotated functions are indicated. Together, these data summarize the depth, coverage, and abundance distribution of the sporozoite proteome.

The most abundant proteins detected in sporozoites include actin, glyceraldehyde-3-phosphate dehydrogenase (GAPDH), thioredoxin peroxidase-like protein, protein disulfide isomerase, two heat shock protein 70 (HSP70) isoforms, type E aldehyde/alcohol dehydrogenase (AdhE), nucleoside diphosphate kinase, heat shock protein 90 (HSP90), and three hypothetical proteins (Fig 1D). These proteins represent diverse functional categories, including cytoskeletal organization, energy metabolism, protein folding, and redox homeostasis.

The identified proteome represents 57.6% of the predicted protein-coding genes based on the original annotation of the *C. parvum* Iowa II strain genome (n = 3,944) and 58.3% based on the reannotated Iowa-ATCC strain genome (n = 3,896) [26,27]. Previous large-scale proteomic studies of *C. parvum* sporozoites reported 1,237 proteins using a multi-platform LC–MS/MS approach [21] or 1,107 proteins using a hyperLOPIT-based spatial proteomics strategy [20], while an oocyst wall–enriched proteome identified 798 proteins [24]. In addition, a proteomic analysis of *C. andersoni* identified 1,586 proteins [25]. In comparison, the present study substantially expands the experimentally validated protein repertoire of *C. parvum* sporozoites and provides the most comprehensive sporozoite proteome reported to date (Table 1).

### Weak correlation between the sporozoite proteome and transcriptomes

To evaluate the relationship between protein abundance and transcript levels in excysted *C. parvum* sporozoites, we compared the sporozoite proteome generated in this study with previously reported transcriptomic and proteomic datasets. Comparison with an earlier sporozoite proteome derived from 2D gel electrophoresis followed by LC–MS/MS analysis revealed a strong and statistically significant positive correlation (Pearson correlation coefficient r = 0.7184; P < 0.0001) (Fig 2A), indicating good overall concordance between independently generated proteomic datasets.

**Figure 2.**
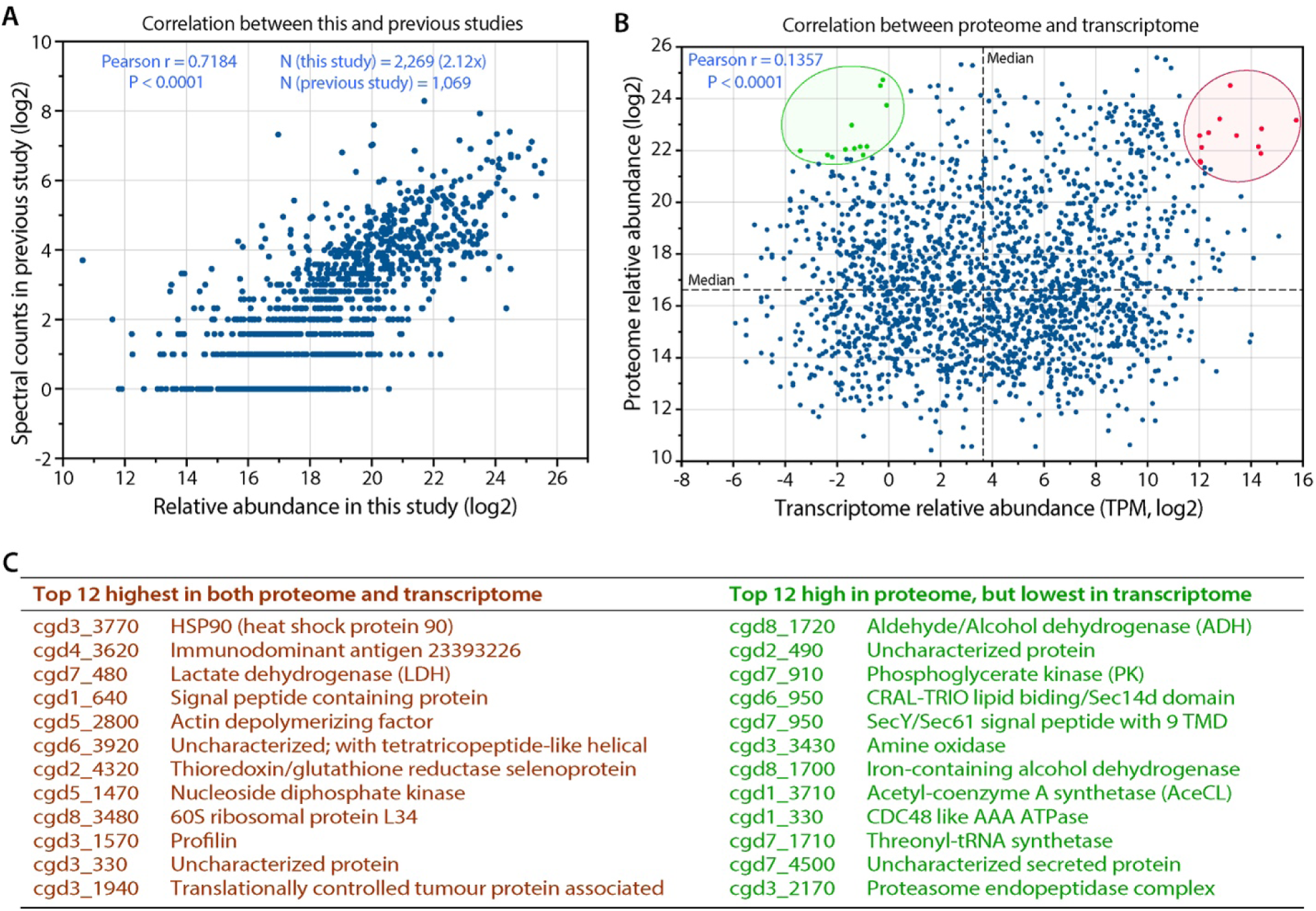
Correlation of the *C. parvum* sporozoite proteome with transcriptomic and previous proteomic datasets. (A) Correlation between protein abundance values obtained in this study and spectral counts reported in a previous sporozoite proteomic analysis. Each point represents an individual protein detected in both datasets. Pearson correlation coefficient (r) and P value are indicated. (B) Correlation between protein abundance values from the sporozoite proteome and transcript abundance values derived from RNA-seq analysis of sporozoites. Protein abundance is shown as log2-transformed proteomic intensity, and transcript abundance is shown as log2-transformed transcripts per million (TPM). Pearson correlation coefficient (r) and P value are indicated. (C) Examples of proteins illustrating the discordance between protein abundance and transcript levels in sporozoites. Proteins with high abundance in both proteome and transcriptome, as well as proteins with high proteomic abundance but low transcript abundance, are highlighted and labeled. Together, these analyses illustrate strong concordance between independent proteomic datasets but weak correspondence between protein abundance and transcript levels in excysted sporozoites.

In contrast, comparison of protein abundance with RNA-seq–based sporozoite transcript levels showed a very weak correlation (Pearson r = 0.1357; P < 0.0001) (Fig 2B). Proteins with high abundance in the proteome were associated with a wide range of transcript abundances, including both highly expressed and lowly expressed transcripts (Fig 2C). This pattern was also observed among enzymes within the same metabolic pathway. For example, in the glycolytic pathway, lactate dehydrogenase (LDH), aldehyde/alcohol dehydrogenase (ADH), phosphoglycerate kinase (PK), and acetyl-CoA synthetase (AceCL) are among the most abundant proteins detected, whereas their corresponding transcripts span from among the highest (LDH) to among the lowest (ADH, PK, and AceCL) abundance groups.

The observed proteome–transcriptome correlation in *C. parvum* sporozoites is markedly lower than the global correlations typically reported for eukaryotes, which generally range from approximately 0.2 to 0.5 [41], and is also substantially lower than correlations reported for *Plasmodium falciparum* sporozoites and other developmental stages (r = 0.370–0.598) [42]. These results indicate that transcript abundance is a poor predictor of protein abundance in excysted *C. parvum* sporozoites.

### Functional composition of the sporozoite proteome highlights enrichment of carbohydrate metabolism among highly abundant proteins

To assess the functional landscape of the excysted *C. parvum* sporozoite proteome, identified proteins were mapped to Kyoto Encyclopedia of Genes and Genomes (KEGG) pathways. Of the 2,272 proteins detected, 876 could be assigned to at least one KEGG pathway (Fig 3, left). The majority of these proteins were classified under categories related to genetic information processing, including “Genetic information processing” (n = 378) and “Protein families: genetic information processing” (n = 194), together accounting for approximately two-thirds (65.3%) of all KEGG-mapped proteins. These categories encompass proteins involved in transcription, translation, protein folding, degradation, and associated processes. Other functional categories contained substantially fewer proteins, including “Cellular processes” (n = 49), “Environmental information processing” (n = 38), and “Carbohydrate metabolism” (n = 28), along with several additional metabolic pathways.

**Figure 3.**
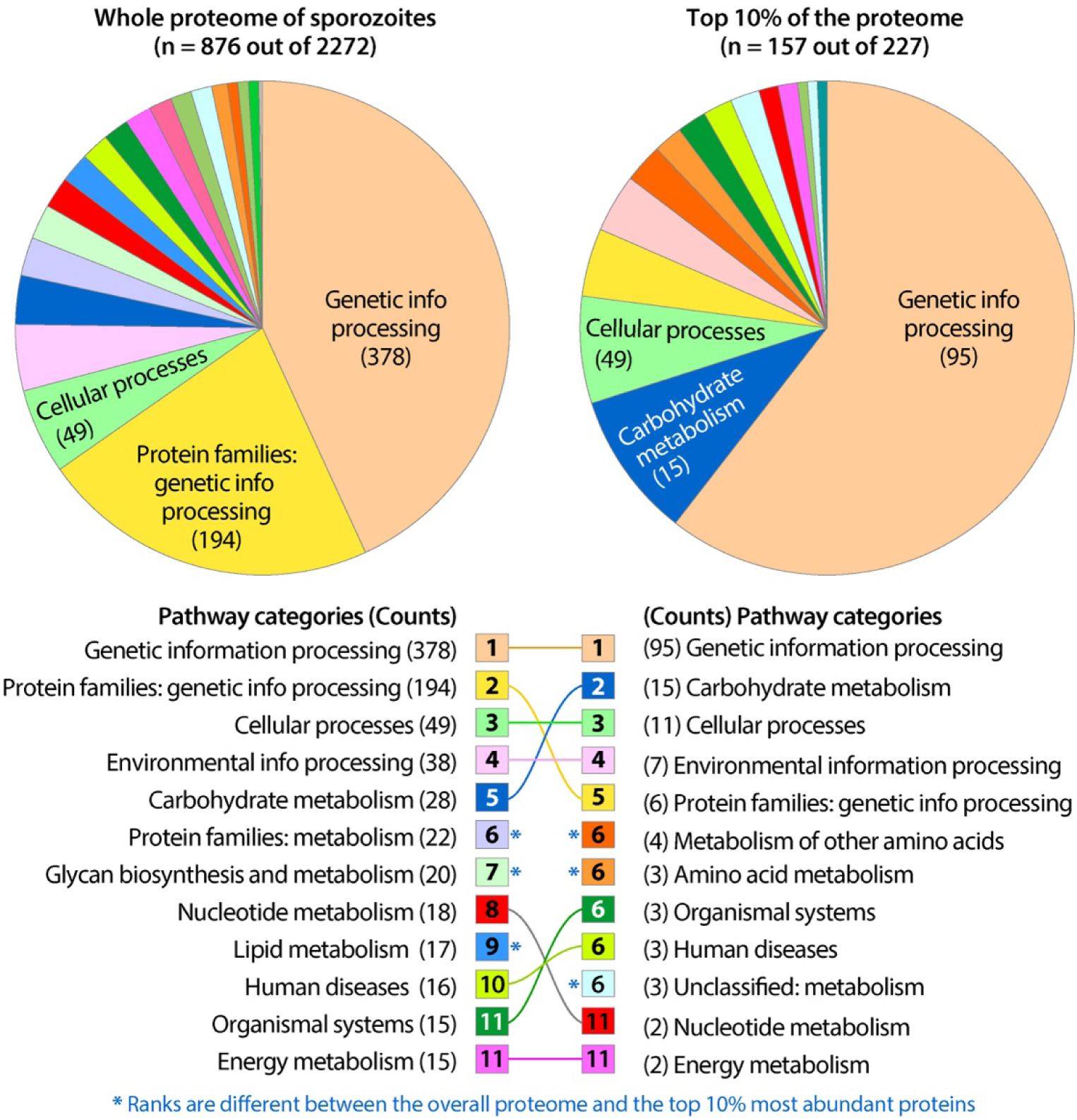
Functional composition of the *C. parvum* sporozoite proteome and abundance-weighted enrichment analysis. Proteins identified in the sporozoite proteome were assigned to Kyoto Encyclopedia of Genes and Genomes (KEGG) functional categories based on KEGG ortholog (KO) annotations. (Left) Distribution of KEGG pathway categories among all KEGG-mapped proteins detected in the sporozoite proteome (n = 876). Numbers in parentheses indicate the number of proteins assigned to each category. (Right) Distribution of KEGG pathway categories among the top 10% most abundant proteins in the proteome (n = 227), of which 157 could be mapped to KEGG pathways. Pathway categories are ranked independently for the complete proteome and the abundance-weighted subset, and asterisks indicate categories whose relative rankings differ between the two analyses. These comparisons illustrate differences between overall functional composition and enrichment among the most abundant proteins in excysted sporozoites.

To further evaluate functional enrichment among the most abundant proteins, we examined the top 10% of proteins ranked by proteomic signal intensity (n = 227). Of these, 157 proteins could be mapped to KEGG pathways (Fig 3, right). As observed for the complete proteome, proteins associated with genetic information processing remained the most abundant functional class (n = 95). Notably, however, proteins involved in carbohydrate metabolism ranked second in this abundance-weighted analysis (n = 15), followed by proteins associated with cellular processes (n = 11). In addition, seven enzymes involved in amino acid–related metabolism were present among the top-ranked categories, including those classified under “Metabolism of other amino acids” (n = 4) and “Amino acid metabolism” (n = 3).

Because *Cryptosporidium* lacks de novo amino acid biosynthesis, enzymes categorized under amino acid metabolism are primarily involved in peripheral reactions or interconversions rather than core synthetic pathways. These include enzymes associated with redox balance, methyl-group metabolism, and amino acid turnover, such as 3-hydroxyacyl-CoA dehydrogenase (HSD17B10), S-adenosylmethionine synthetase (SAMS), adenosylhomocysteinase (AHCY), glutathione peroxidase (GPx), leucyl aminopeptidase (PepA), aminopeptidase N (PepN), and NADP-dependent thioredoxin reductase (TXNRD). Together, these analyses indicate that while proteins involved in genetic information processing dominate the sporozoite proteome overall, carbohydrate metabolism is disproportionately represented among the most abundant proteins detected in excysted sporozoites.

### Global features of the sporozoite phosphoproteome and phosphorylation motifs

Phosphoproteomic analysis of excysted *C. parvum* sporozoites identified a total of 8,994 phosphorylation sites distributed across 5,690 phosphopeptides and mapped to 2,141 proteins (Fig 4A; Table S2). Phosphopeptide lengths ranged from 7 to more than 26 amino acids, with the majority clustering between 11 and 15 residues (Fig 4B). Among all identified phosphorylation sites, serine phosphorylation was the most prevalent (p-Ser; n = 6,703; 74.53%), followed by threonine phosphorylation (p-Thr; n = 2,179; 24.23%) and tyrosine phosphorylation (p-Tyr; n = 112; 1.25%) (Fig 4C).

**Figure 4.**
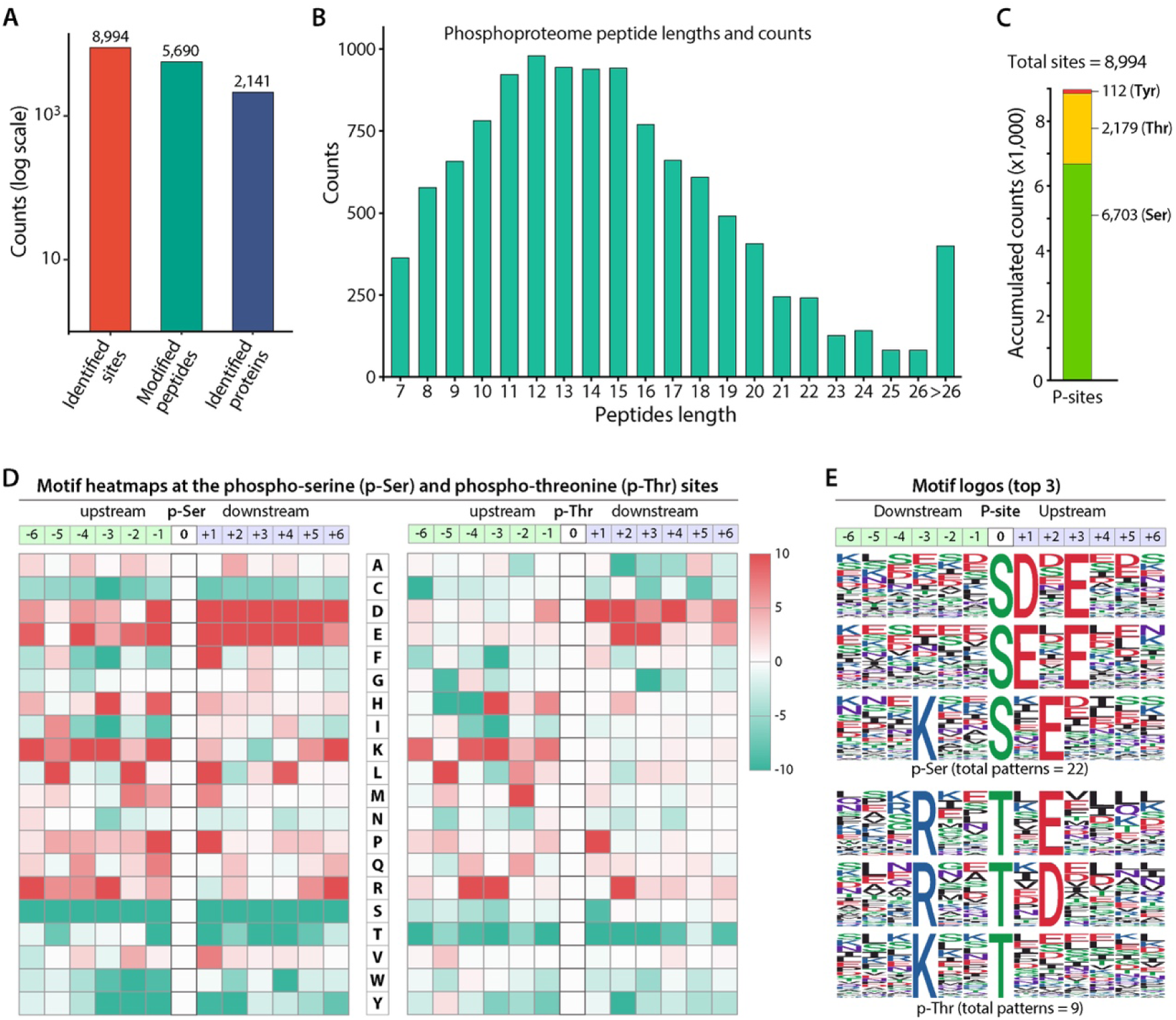
Global features of the *C. parvum* sporozoite phosphoproteome and phosphorylation motifs. (A) Summary of the phosphoproteomic dataset, showing the numbers of identified phosphorylation sites, modified peptides, and phosphoproteins detected in excysted sporozoites. (B) Length distribution of identified phosphopeptides. (C) Distribution of phosphorylation site types, showing the numbers of serine (p-Ser), threonine (p-Thr), and tyrosine (p-Tyr) phosphorylation sites detected. (D) Heatmaps depicting amino acid frequency enrichment and depletion in the upstream and downstream regions flanking p-Ser and p-Thr sites. Amino acid positions are shown relative to the phosphorylated residue (position 0). (E) Sequence logos representing the top three statistically significant phosphorylation motifs for p-Ser and p-Thr sites, identified by motif analysis. Together, these panels summarize the composition, residue distribution, and sequence features of phosphorylation sites identified in excysted *C. parvum* sporozoites.

Sequence motif analysis of serine- and threonine-phosphorylated peptides revealed statistically significant enrichment of specific amino acid patterns flanking the modified residues. For both p-Ser and p-Thr sites, acidic residues (aspartic acid and glutamic acid) were enriched in downstream positions, whereas basic residues (lysine and arginine) were more frequently observed upstream of the phosphorylation sites (Fig 4D). In contrast, neutral residues such as cysteine, serine, threonine, tryptophan, and tyrosine were underrepresented in the flanking regions. Motif-x analysis identified a total of 31 significant motif patterns, including 22 motifs centered on p-Ser sites and 9 motifs centered on p-Thr sites. The top three motifs for each phosphorylation type, ranked by motif score, are shown as sequence logos in Fig 4E, and the complete list of identified motifs and associated statistical parameters is provided in Table S3.

The overall distribution of phosphorylation site types in *C. parvum* sporozoites is similar to those reported in other eukaryotic systems, including humans, yeast, and the apicomplexan parasite *Plasmodium falciparum*, in which serine phosphorylation predominates, followed by threonine and a minor fraction of tyrosine phosphorylation [43–46].

### Immunofluorescence analysis reveals distinct spatial distributions of phosphorylated proteins in sporozoites

To independently visualize protein phosphorylation in excysted *C. parvum* sporozoites, immunofluorescence assays (IFA) were performed using pan-phosphoprotein antibodies recognizing phospho-serine (p-Ser), phospho-threonine (p-Thr), or phospho-tyrosine (p-Tyr) residues (Fig 5). Under identical labeling and imaging conditions, strong fluorescence signals were observed for p-Ser and p-Thr, whereas p-Tyr signals were consistently weaker (Fig 5A–C), consistent with the relative frequencies of phosphorylation site types identified in the phosphoproteomic dataset.

**Figure 5.**
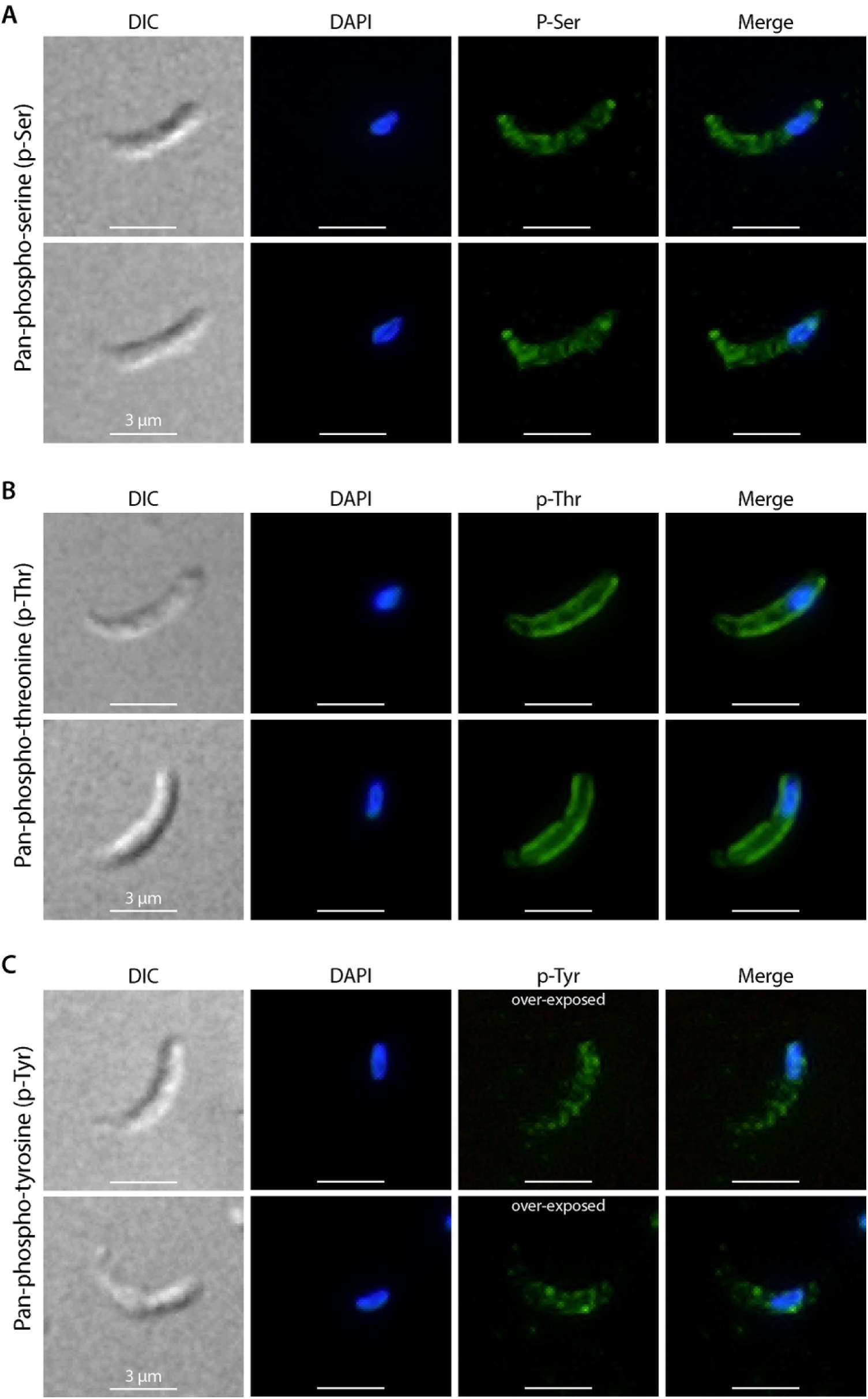
Immunofluorescence visualization of phosphorylated proteins in excysted *C. parvum* sporozoites. Immunofluorescence assays were performed on excysted sporozoites using pan-phosphoprotein antibodies recognizing phospho-serine (p-Ser), phospho-threonine (p-Thr), or phospho-tyrosine (p-Tyr) residues. (A) Sporozoites labeled with anti-p-Ser antibody. (B) Sporozoites labeled with anti-p-Thr antibody. (C) Sporozoites labeled with anti-p-Tyr antibody. For each panel, differential interference contrast (DIC), DAPI nuclear staining, phosphoprotein signal, and merged images are shown. Scale bars, 3 μm. Strong p-Ser and p-Thr signals and comparatively weaker p-Tyr signals are observed under identical labeling and imaging conditions. Distinct spatial distribution patterns are evident for different classes of phosphorylated proteins, providing independent experimental support for the phosphoproteomic dataset.

Labeling with the anti–p-Ser antibody produced broadly distributed fluorescence throughout the sporozoite cytoplasm, often with a granular appearance and slightly enhanced signal intensity toward both ends of the parasite (Fig 5A). A similar, though weaker, overall distribution pattern was observed with the anti–p-Tyr antibody (Fig 5C), indicating that the majority of p-Ser– and p-Tyr–containing proteins are broadly distributed within the sporozoite body. In contrast, labeling with the anti–p-Thr antibody resulted in a strong peripheral signal outlining the sporozoite (Fig 5B). The p-Thr fluorescence was largely confined within the parasite boundary and did not extend beyond the cell edge, consistent with localization to internal membrane-associated structures.

Together, these observations confirm the presence of serine, threonine, and tyrosine phosphorylation in excysted *C. parvum* sporozoites and demonstrate distinct spatial distribution patterns for different classes of phosphorylated proteins, providing independent experimental support for the phosphoproteomic findings.

### Comparative analysis of the sporozoite proteome and phosphoproteome reveals pathway-level differences in phosphorylation representation

The phosphoproteomic dataset comprised 5,690 phosphopeptides with signal intensities spanning a wide dynamic range, from 3.29 to 123,596.32 (mean = 1,350.36; median = 311.13) (Table S2). To reduce potential contributions from low-intensity signals and nonspecific peptide binding during immobilized metal affinity chromatography enrichment, phosphopeptides with signal intensities below 400 were excluded from subsequent analyses. The filtered dataset mapped to 833 phosphorylated proteins, representing 36.9% of the proteins detected in the sporozoite proteome (Fig 6A; Table S4).

**Figure 6.**
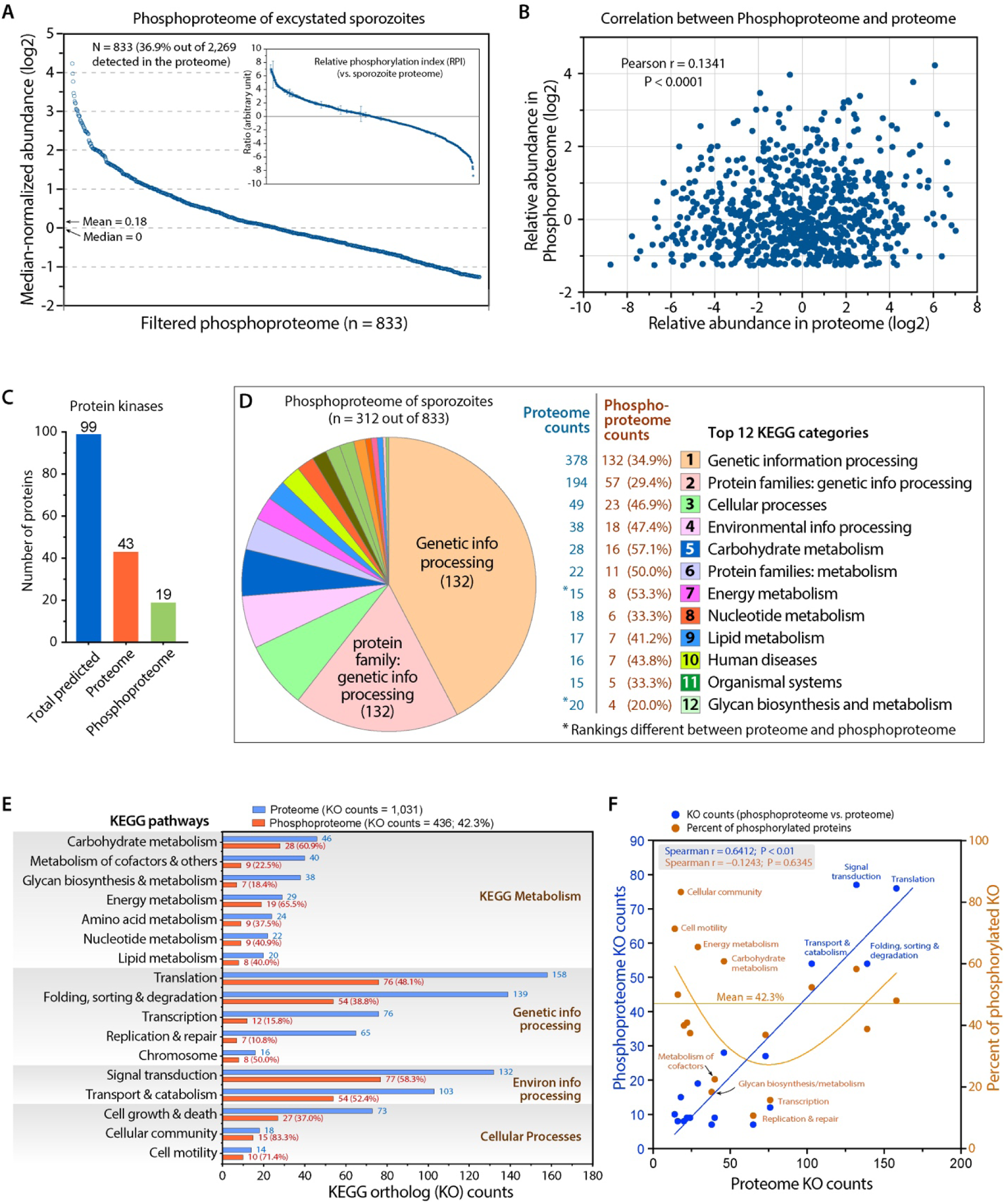
Comparative analysis of the *C. parvum* sporozoite proteome and phosphoproteome. (A) Distribution of median-normalized phosphoprotein abundance values (log2) for the filtered phosphoproteome (n = 833 proteins). The inset shows the distribution of relative phosphorylation index (RPI) values derived from comparison of phosphoproteome and proteome datasets. (B) Correlation between protein abundance in the sporozoite proteome and phosphoprotein abundance in the phosphoproteome. Each point represents an individual protein detected in both datasets. Pearson correlation coefficient (r) and P value are indicated. (C) Detection of predicted protein kinases in the sporozoite proteome and phosphoproteome. Bars indicate the number of kinases identified in each dataset relative to the total number of kinases predicted from the genome. (D) Distribution of Kyoto Encyclopedia of Genes and Genomes (KEGG) ortholog (KO) counts across major functional pathway categories in the sporozoite proteome and phosphoproteome. (E) Comparison of KO counts between proteome and phosphoproteome datasets across KEGG pathway categories. (F) Relationship between KO counts in the proteome and phosphoproteome (blue) and the percentage of phosphorylated proteins within each pathway (brown). Spearman correlation coefficient and P value are indicated. Together, these analyses illustrate pathway-level differences in phosphorylation representation that are not directly proportional to protein abundance in excysted sporozoites.

Comparison of protein abundance between the phosphoproteome and proteome revealed a weak overall correlation (Pearson r = 0.1341; P < 0.0001) (Fig 6B), indicating that phosphorylation-derived signal intensity is largely independent of protein abundance. Of the 99 protein kinases predicted in the *C. parvum* genome, 43 (43.4%) were detected in the sporozoite proteome, and 19 of these were also identified in the phosphoproteome (Fig 6C), demonstrating that a subset of expressed kinases is itself phosphorylated in excysted sporozoites.

Functional mapping of the 833 phosphorylated proteins to KEGG pathways assigned 312 proteins to defined pathways (Fig 6D and 6E). The overall distribution of KEGG ortholog (KO) counts across pathways in the phosphoproteome broadly mirrored that observed in the proteome, with “Genetic information processing,” “Protein families: genetic information processing,” “Cellular processes,” “Environmental information processing,” and “Carbohydrate metabolism” ranking among the most represented categories in both datasets. However, the relative proportion of phosphorylated proteins varied substantially between pathways. While KO counts between proteome and phosphoproteome datasets were moderately correlated across pathways (Spearman r = 0.6412; P < 0.01) (Fig 6F, blue), the fraction of phosphorylated proteins within individual pathways showed marked differences (Fig 6F, brown).

Pathways with lower overall KO representation, including those associated with cellular community, cell motility, energy metabolism, and carbohydrate metabolism, exhibited comparatively higher proportions of phosphorylated proteins. In contrast, pathways with higher KO counts, such as transcription, replication and repair, glycan biosynthesis and metabolism, and metabolism of cofactors, showed lower relative phosphorylation representation. Together, these analyses demonstrate that phosphorylation is unevenly distributed across functional pathways in excysted *C. parvum* sporozoites and highlight pathway-level differences in phosphorylation representation that are not directly proportional to protein abundance.

### Proteome, phosphoproteome, and relative phosphorylation index profiles of glycolytic and connected enzymes

To examine phosphorylation patterns within carbohydrate metabolism in greater detail, we analyzed the abundance of enzymes involved in glycolysis and major connected pathways using three complementary measures: protein abundance in the proteome, phosphoprotein abundance in the phosphoproteome, and relative phosphorylation index (RPI) derived from the comparison of the two datasets (Fig 7; also see Fig 6A inset for the RPI distribution curve). A total of 40 enzymes (including isozymes) participating in 35 reactions were annotated across glycolysis and associated metabolic branches (Fig 7A–C; Table S5).

**Figure 7.**
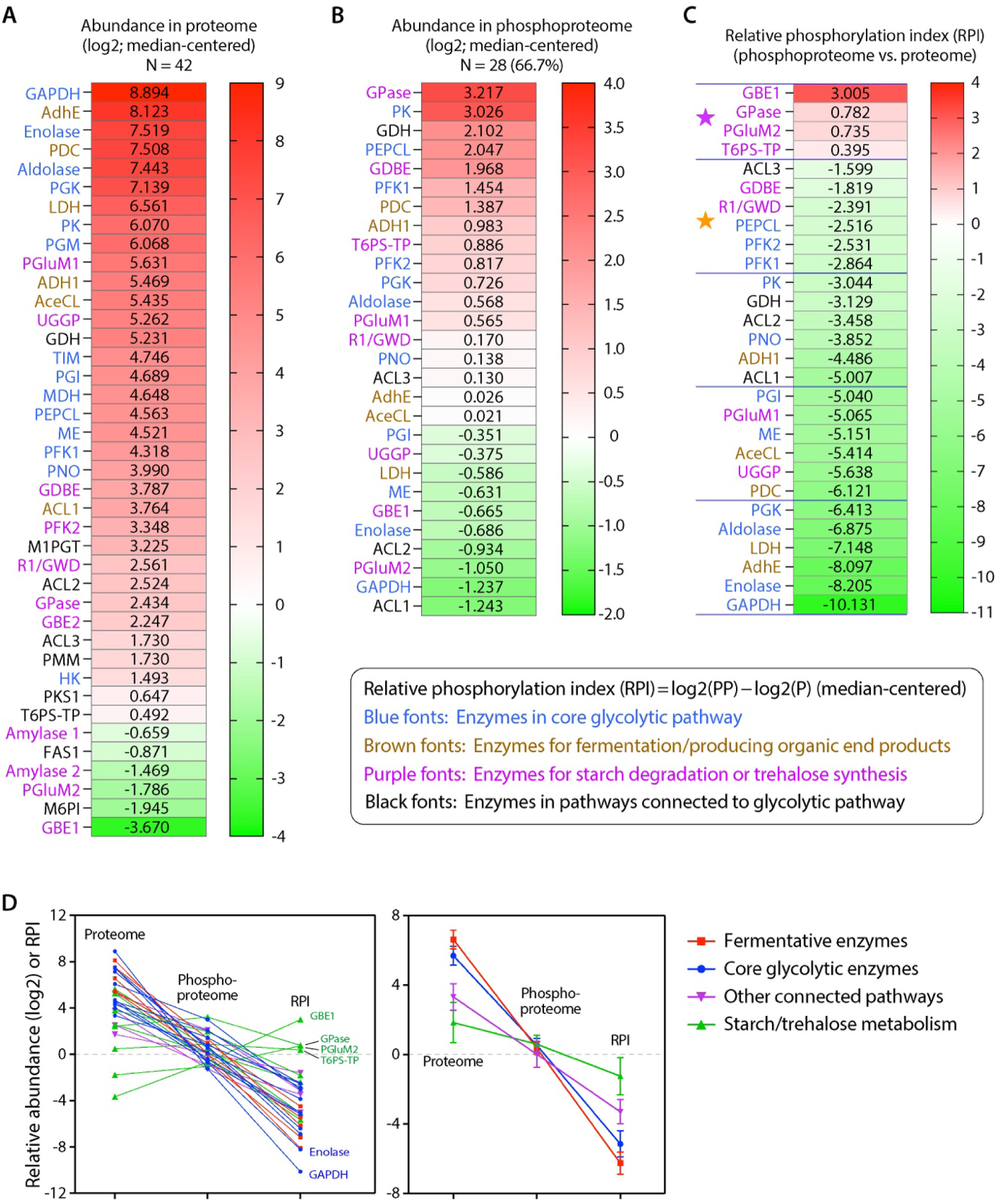
Proteome abundance, phosphoproteome abundance, and relative phosphorylation index of glycolytic and connected enzymes in excysted *C. parvum* sporozoites. (A) Heatmap showing median-centered protein abundance (log2) of enzymes involved in glycolysis and major connected pathways, based on proteomic data. Enzymes are grouped by pathway affiliation, including core glycolysis, fermentation/organic end product formation, starch degradation, and trehalose metabolism. (B) Heatmap showing median-centered phosphoprotein abundance (log2) for the same set of enzymes detected in the phosphoproteome. Enzymes not detected in the phosphoproteome are indicated accordingly. (C) Relative phosphorylation index (RPI) values for enzymes detected in both proteome and phosphoproteome datasets. RPI values were calculated as the difference between median-centered phosphoproteome and proteome abundance values and are shown on a log2 scale. Top 10 enzymes by RPI were marked with stars, with top four with above-median RPI in purple stars, and the following six in orange stars. (D) Comparison of enzyme protein abundance and RPI values across the pathway, illustrating the relationship between overall protein expression and phosphorylation propensity. Enzyme names and corresponding gene identifiers are indicated, and color scales are shown for each panel. **Abbreviations:** AceCL, acetyl-coenzyme A synthetase; ACL, fatty acyl-CoA synthetase; AdhE, acetaldehyde/alcohol dehydrogenase (type E); ADH1, alcohol dehydrogenase (monofunctional); FAS1, fatty acid synthase (type I); GAPDH, glyceraldehyde 3-phosphate dehydrogenase; GBE1, glycogen/glucan branching enzyme; GBE2, glycogen/glucan branching enzyme; GDBE, glycogen debranching enzyme; GDH, glycerol-3-phosphate dehydrogenase; GPase, glycogen phosphorylase; HK, hexokinase; LDH, lactate dehydrogenase; M1PGT, mannose-1-phosphate guanylyltransferase; M6PI, mannose-6-phosphate isomerase; MDH, malate dehydrogenase; ME, malic enzyme; PDC, pyruvate decarboxylase; PEPCL, phosphoenolpyruvate carboxylase; PGI, phosphoglucose isomerase; PGK, phosphoglycerate kinase; PGluM, phosphoglucomutase; PK, pyruvate kinase; PKS1, polyketide synthase (type I); PMM, phosphomannomutase; PNO, Pyruvate:NADP+ oxidoreductase; PPi-PFK, PPi-dependent phosphofructokinase; R1/GWDK, R1-like alpha-glucan water dikinase; T6PS-TP, trehalose-6-phosphate synthase; TIM, triosephosphate isomerase; UGGP, UDP-N-acetylglucosamine pyrophosphorylase.

Proteomic analysis detected 39 of the annotated enzymes, with the exception of acetyl-CoA carboxylase (ACC), which was not detected in the sporozoite proteome (Fig 7A). Most enzymes exhibited above-median protein abundance (n = 36; 85.7%), consistent with the prominence of carbohydrate metabolism among highly abundant proteins. Core glycolytic and fermentative enzymes, including glyceraldehyde-3-phosphate dehydrogenase (GAPDH), enolase, aldolase, pyruvate decarboxylase (PDC), and type E aldehyde/alcohol dehydrogenase (AdhE), ranked among the most abundant proteins detected. In contrast, several enzymes involved in starch degradation or peripheral pathways, such as glucan branching enzyme isoform 1 (GBE1), mannose-6-phosphate isomerase (M6PI), phosphoglucomutase isoform 2 (PGluM2), and α-amylase isoform 2, exhibited comparatively low protein abundance.

Phosphoproteomic analysis identified 28 enzymes within the same pathway set (Fig 7B). Among these, 18 enzymes (64.3%) displayed above-median phosphoprotein abundance. Enzymes associated with starch metabolism and related processes, including glycogen/glucan phosphorylase (GPase), glucan debranching enzyme (GDBE), and trehalose-6-phosphate synthase–trehalose phosphatase (T6PS-TP), were prominent within the phosphoproteome, whereas several highly abundant proteomic enzymes, such as GAPDH and enolase, exhibited comparatively low phosphoprotein signal. Notably, hexokinase (HK) was not detected in the phosphoproteome, in both filtered and unfiltered datasets including low signal phosphopeptides.

To enable comparative assessment of phosphorylation propensity relative to protein abundance, RPI values were calculated for enzymes detected in both datasets (Fig 7C). Enzymes involved in starch and trehalose metabolism exhibited the highest RPI values, with GBE1, GPase, PGluM2, and T6PS-TP ranking among the top entries. Additional enzymes with elevated RPI values included acetyl-CoA ligase isoform 3 (ACL3), GDBE, glucan-water dikinase (R1/GWD), phosphoenolpyruvate carboxylase (PEPCL), and the two pyrophosphate-dependent phosphofructokinase isozymes (PFK1 and PFK2). In contrast, several highly abundant glycolytic enzymes, including GAPDH, enolase, and aldolase, displayed low RPI values. Across the pathway, enzyme protein abundance and RPI showed an inverse relationship, with enzymes of high proteomic abundance generally exhibiting lower phosphorylation propensity and vice versa (Fig 7D).

### Integrated reconstruction of glycolysis and connected metabolic pathways in excysted sporozoites

To provide a pathway-level overview of carbohydrate metabolism in excysted *C. parvum* sporozoites, proteome and phosphoproteome data were integrated into a reconstructed glycolytic map with major connected metabolic branches (Fig 8; also see Fig S1 for a simplified pathway schematic). The reconstruction includes enzymes involved in glycolysis, fermentation, starch degradation, trehalose metabolism, and selected auxiliary reactions, with median-centered protein abundance, phosphoprotein abundance, and relative phosphorylation index (RPI) values displayed for each detected enzyme.

**Figure 8.**
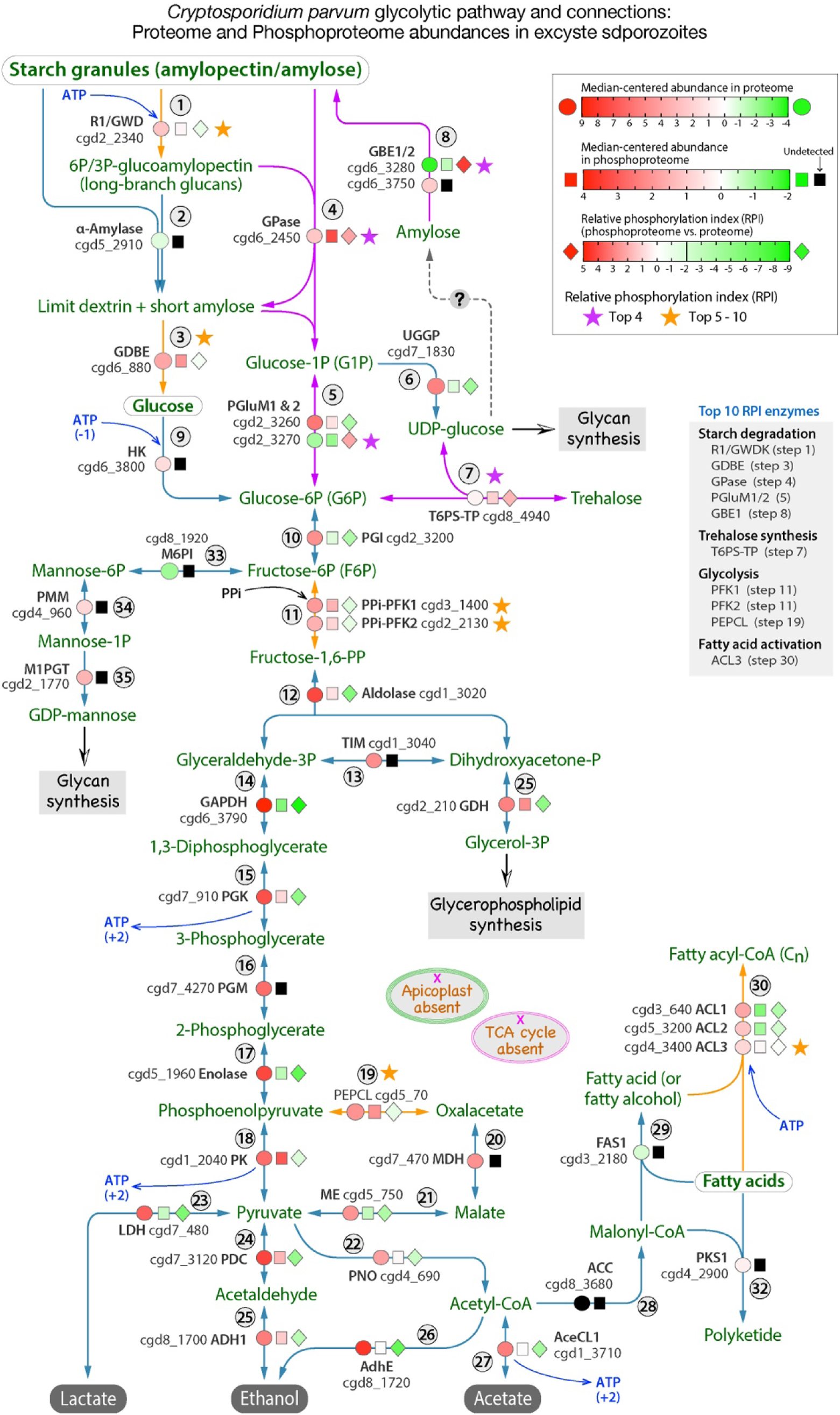
Integrated reconstruction of glycolysis and connected carbohydrate metabolic pathways in excysted Cryptosporidium parvum sporozoites. Proteome and phosphoproteome data were integrated into a reconstructed pathway map encompassing glycolysis, fermentation, starch degradation, trehalose metabolism, and selected connected reactions. Enzymes are annotated with median-centered protein abundance values derived from the proteome (log2 scale), median-centered phosphoprotein abundance values derived from the phosphoproteome (log2 scale), and relative phosphorylation index (RPI) values calculated from the comparison of the two datasets. Detected enzymes are color-coded according to abundance or RPI as indicated by the legends, whereas undetected enzymes are marked accordingly. Core glycolytic reactions, branch points connecting storage carbohydrate metabolism to glycolysis, and downstream fermentative steps are indicated. This integrated representation summarizes coverage, abundance, and phosphorylation propensity of carbohydrate metabolic enzymes in excysted sporozoites and provides a pathway-level framework for visualizing proteome and phosphoproteome data. Abbreviations refer to Fig 7.

Proteomic analysis revealed near-complete coverage of the glycolytic pathway, with all core enzymes detected except for acetyl-CoA carboxylase (ACC). Enzymes catalyzing central glycolytic steps, including glyceraldehyde-3-phosphate dehydrogenase (GAPDH), enolase, aldolase, phosphoglycerate kinase (PGK), phosphoglycerate mutase (PGM), and pyruvate kinase (PK), exhibited high protein abundance across the pathway. In contrast, enzymes involved in starch degradation and trehalose synthesis, such as glucan branching enzymes (GBE1/2), glucan phosphorylase (GPase), glucan debranching enzyme (GDBE), glucan-water dikinase (R1/GWD), and trehalose-6-phosphate synthase–trehalose phosphatase (T6PS-TP), were detected at lower protein abundance levels.

Phosphoproteomic data demonstrated heterogeneous phosphorylation patterns across the reconstructed network. Multiple enzymes in starch and trehalose metabolism displayed high phosphoprotein abundance and elevated RPI values, whereas several highly abundant glycolytic enzymes exhibited comparatively low phosphorylation propensity. Enzymes ranking among the highest by RPI included GBE1, GPase, PGluM2, T6PS-TP, ACL3, and R1/GWD, while GAPDH, enolase, aldolase, and LDH ranked among the lowest. Notably, several enzymes with elevated RPI values catalyze steps upstream of glycolysis or branch points connecting glycolysis to storage carbohydrate metabolism.

The integrated pathway reconstruction highlights distinct abundance and phosphorylation profiles among enzymes participating in carbohydrate metabolism in excysted *C. parvum* sporozoites and provides a consolidated framework for visualizing proteome and phosphoproteome data within a metabolic context.

## Discussion

In this study, we present an integrated proteomic and phosphoproteomic analysis of excysted *Cryptosporidium parvum* sporozoites, providing the most comprehensive proteome reported for this invasive stage and establishing the first global phosphoproteome for the parasite. By combining quantitative proteomics, phosphopeptide enrichment, motif analysis, and immunofluorescence validation, we define both the protein expression landscape and the distribution of phosphorylation events in sporozoites. Several overarching features emerge from these analyses. First, protein abundance in sporozoites shows only a weak relationship with transcript levels, underscoring extensive post-transcriptional regulation at this stage. Second, protein phosphorylation is widespread but unevenly distributed across functional pathways, with a notable enrichment among enzymes involved in carbohydrate metabolism. Finally, comparative analysis using a relative phosphorylation index highlights distinct phosphorylation propensity among metabolic enzymes that is not simply proportional to protein abundance. Together, these findings provide new insights into post-transcriptional and post-translational regulation in *C. parvum* sporozoites and establish a resource for future studies of parasite metabolism and signaling.

A striking feature of the excysted *C. parvum* sporozoite dataset is the very weak correlation between protein abundance and transcript levels. While independent proteomic datasets show strong concordance with one another, protein abundance correlates poorly with RNA-seq–derived transcript abundance (Pearson r ≈ 0.14), indicating that transcript levels are a poor predictor of protein expression in this invasive stage. This degree of discordance is substantially lower than that typically observed in eukaryotic systems, where global proteome–transcriptome correlations generally range from 0.2 to 0.5 [41], and is also lower than values reported for *Plasmodium falciparum* across multiple developmental stages, including sporozoites (r = 0.370–0.598) [42].

Such pronounced uncoupling between transcript and protein abundance suggests extensive post-transcriptional regulation in *C. parvum* sporozoites. This phenomenon is consistent with the biology of excystation and host cell invasion, processes that require rapid deployment of pre-existing protein machinery rather than de novo transcription. In this context, transcripts detected in sporozoites may reflect transcriptional activity during earlier developmental stages, while protein abundance more directly represents the functional state of the parasite at the time of host cell contact. Similar reliance on translational control and mRNA storage has been described in other apicomplexan parasites, particularly during transitions between extracellular and intracellular stages, although the magnitude of proteome–transcriptome discordance observed here appears especially pronounced.

The weak correlation is also evident within individual metabolic pathways, where enzymes with comparable functional roles exhibit widely divergent transcript abundances despite similar or even higher protein abundance. For example, several core glycolytic and fermentative enzymes are among the most abundant proteins detected in sporozoites, yet their corresponding transcripts span from among the highest to among the lowest abundance groups. Together, these observations underscore the importance of direct protein-level measurements for understanding the biology of excysted *C. parvum* sporozoites and caution against inferring functional activity solely from transcriptomic data in this stage.

Beyond global protein expression, this study provides the first systems-level view of protein phosphorylation in *C. parvum* sporozoites. The phosphoproteome is characterized by a predominance of serine phosphorylation, followed by threonine and a minor fraction of tyrosine phosphorylation, a distribution consistent with that reported in other eukaryotic organisms and apicomplexan parasites [43–46]. Motif analysis further revealed conserved sequence features surrounding phosphorylated residues, including enrichment of acidic residues downstream and basic residues upstream of modification sites, suggesting the involvement of kinase activities with preferences broadly similar to those described in other eukaryotic systems. Together with immunofluorescence validation, these observations indicate that phosphorylation is widespread in excysted sporozoites and exhibits structured, non-random patterns.

At the pathway level, phosphorylation is unevenly distributed across functional categories. While proteins involved in genetic information processing dominate the sporozoite proteome overall, a comparatively higher fraction of proteins associated with carbohydrate metabolism, energy metabolism, cell motility, and cellular community pathways are phosphorylated. This pattern contrasts with pathways such as transcription, replication and repair, and glycan biosynthesis, which show relatively lower proportions of phosphorylated proteins despite substantial representation in the proteome. These differences suggest that phosphorylation in sporozoites preferentially targets specific functional modules rather than reflecting protein abundance alone.

The pathway-level skew toward metabolic processes is particularly notable given the extracellular and invasive nature of the sporozoite stage. Sporozoites must rapidly adapt to changing environmental conditions during excystation and host cell contact, processes that are likely to place strong demands on energy production and metabolic flux. In this context, phosphorylation may provide a flexible means of modulating metabolic enzyme activity, protein–protein interactions, or subcellular localization without requiring new protein synthesis. Although the present data do not establish causal roles for individual phosphorylation events, the enrichment of phosphorylated proteins within defined metabolic pathways highlights phosphorylation as a prominent layer of post-translational organization in *C. parvum* sporozoites.

The pathway-level analyses further reveal a pronounced contrast between protein abundance and phosphorylation propensity within carbohydrate metabolism. Core glycolytic enzymes, including glyceraldehyde-3-phosphate dehydrogenase, enolase, aldolase, phosphoglycerate kinase, and pyruvate kinase, are among the most abundant proteins detected in excysted sporozoites but generally display low relative phosphorylation propensity. In contrast, enzymes involved in starch degradation, trehalose metabolism, and branch points connecting storage carbohydrates to glycolysis exhibit comparatively low protein abundance yet rank among the most highly phosphorylated proteins. This inverse relationship between abundance and phosphorylation propensity suggests that phosphorylation is not uniformly applied across the pathway but instead preferentially associated with specific metabolic nodes.

Notably, many enzymes with elevated phosphorylation propensity catalyze reactions upstream of glycolysis or at branch points linking glycolysis to storage carbohydrate metabolism, including glucan branching and debranching enzymes, glucan phosphorylase, trehalose-6-phosphate synthase–trehalose phosphatase, and phosphoglucomutase isoforms. These enzymes are positioned at interfaces between polysaccharide storage, sugar mobilization, and central carbon metabolism, where modulation of enzymatic activity or interaction states could influence the availability of glycolytic substrates. In contrast, the consistently high abundance and low phosphorylation propensity of core glycolytic enzymes may reflect a constitutively active metabolic backbone that supports the energetic demands of excystation and host cell invasion.

The metabolic architecture of *Cryptosporidium* is characterized by streamlined biosynthetic capacity and a heavy reliance on carbohydrate utilization, fermentation, and salvage pathways. Within this context, differential phosphorylation of enzymes associated with storage carbohydrate turnover may provide a means to fine-tune carbon flux during rapid environmental transitions encountered by sporozoites. Although the present data do not establish direct regulatory effects of phosphorylation on enzyme activity, the observed patterns point to phosphorylation as a potential contributor to metabolic flexibility in the invasive stage of *C. parvum*.

To facilitate comparison between independently acquired proteome and phosphoproteome datasets, we introduced the relative phosphorylation index (RPI) as a normalized measure of phosphorylation propensity relative to protein abundance. This index is intended as a comparative metric rather than a direct estimate of phosphorylation stoichiometry. By accounting for protein abundance, RPI enables identification of proteins and pathways that are disproportionately represented in the phosphoproteome relative to their overall expression levels, thereby highlighting patterns that may be obscured when considering proteome or phosphoproteome data alone.

A notable example of parasite–host divergence in phosphorylation-based metabolic regulation is hexokinase (HK). In excysted C. parvum sporozoites, HK is among the most abundant glycolytic enzymes detected in the proteome, yet it was not identified in the phosphoproteome, resulting in a low relative phosphorylation index. This contrasts with mammalian systems, in which HK isoforms are frequently phosphorylated and subject to extensive post-translational regulation, with phosphorylation implicated in controlling enzymatic activity, subcellular localization, and integration with signaling pathways [47]. Such differences suggest that *C. parvum* may rely on a more streamlined or constitutively active glycolytic framework during the sporozoite stage, with reduced dependence on phosphorylation-based fine-tuning of key entry-point enzymes.

Importantly, the absence of detectable HK phosphorylation in this dataset does not exclude alternative regulatory mechanisms, including allosteric control, regulation at the level of protein abundance, or other PTMs. Nonetheless, this contrast highlights a fundamental divergence in metabolic regulation between the parasite and its mammalian hosts and underscores the value of phosphoproteomic profiling for revealing parasite-specific regulatory logic that may be exploitable for therapeutic intervention.

Several limitations of this approach should be considered. Phosphoproteomic measurements are influenced by peptide detectability, enrichment efficiency, ionization properties, and the dynamic range of mass spectrometric detection. In addition, the absence of experimentally determined phosphorylation stoichiometry—that is, the fraction of protein molecules modified at a given site or across sites—precludes direct calibration of RPI values to absolute proportions of phosphorylated protein for individual targets. Accurate estimation of such stoichiometry would require targeted quantitative approaches capable of distinguishing phosphorylated and non-phosphorylated peptide populations. Filtering steps applied to reduce low-intensity signals may further bias detection toward more readily enriched phosphopeptides. Consequently, RPI values should be interpreted as relative indicators of phosphorylation enrichment rather than as measures of functional importance.

Despite these caveats, the application of RPI at the pathway and enzyme-class levels reveals consistent and biologically coherent patterns, particularly within carbohydrate metabolism.

Enzymes associated with storage carbohydrate turnover and metabolic branch points exhibit elevated RPI values across the pathway, whereas core glycolytic enzymes show low phosphorylation propensity despite high protein abundance. The concordance of these trends across multiple analytical layers supports the utility of RPI as an exploratory tool for identifying phosphorylation-enriched functional modules and generating hypotheses for targeted functional studies.

Taken together, this study establishes a comprehensive proteomic and phosphoproteomic framework for excysted *Cryptosporidium parvum* sporozoites and provides a foundation for future functional investigations. By defining the largest sporozoite proteome to date and the first global phosphoproteome for this parasite, the dataset offers a resource for exploring post-transcriptional and post-translational regulation in an experimentally challenging organism. The observed pathway-level enrichment of phosphorylation, particularly within carbohydrate metabolism, highlights candidate enzymes and metabolic nodes for targeted validation using genetic, biochemical, or chemical perturbation approaches. Future studies combining quantitative phosphoproteomics with measurements of phosphorylation stoichiometry, kinase–substrate relationships, and temporal dynamics during excystation and host cell invasion will be essential for elucidating the functional consequences of individual phosphorylation events. Beyond advancing our understanding of *Cryptosporidium* biology, the integrative framework and datasets presented here may also inform comparative studies across apicomplexan parasites and support the identification of metabolic vulnerabilities relevant to therapeutic development.

## Acknowledgements

This research was supported by grants of the National Key Research and Development Program of China (2022YFD1800200 to G.Z. and J.Y.).

## Supporting Information

**S1 Fig. Simplified reconstruction of carbohydrate metabolism in excysted *C. parvum* sporozoites.** This schematic summarizes the major glycolytic and connected carbohydrate metabolic pathways identified in this study. Enzymes and pathways are shown for conceptual clarity and do not reflect quantitative protein abundance or phosphorylation levels, which are presented in **Figures 7 and 8**. (PDF)

**S1 Table. Abundance of excysted sporozoite proteome mapped to GenBank accession numbers and CryptoDB gene IDs.** (XLSX)

**S2 Table. Phosphorylation sites in peptides and mass spectrum signal intensities.** (XLSX)

**S3 Table. Motif logos at the Ser/S and Thr/T phosphorylated sites in Cryptosporidium parvum sporozoites and statistical parameters based on motif-x algorithm.** (XLSX)

**S4 Table. Merged phosphoproteome and proteome datasets with abundances (log2 values) and protein IDs and descriptions.** In proteome dataset, peptides with intensities lower than 400 were filtered to reduce potential contributions from low-intensity signals and nonspecific peptide binding during immobilized metal affinity chromatography enrichment. (XLSX)

**S5 Table. Enzymes in the glycolytic and connected pathways: proteome and phosphoproteome abundances, and relative phosphorylation index (RPI).** (XLSX)

## Author Contributions

**Conceptualization:** Guan Zhu.

**Data curation:** Haitao Li, Dongqiang Wang, Guan Zhu.

**Formal analysis:** Dongqiang Wang, Meng Li, Guan Zhu.

**Funding acquisition:** Jigang Yin, Guan Zhu.

**Investigation:** Dongqiang Wang, Meng Li, Chenchen Wang.

**Methodology:** Dongqiang Wang, Meng Li, Guan Zhu.

**Project administration:** Guan Zhu.

**Supervision:** Jigang Yin, Guan Zhu.

**Validation:** Guan Zhu.

**Visualization:** Haitao Li, Guan Zhu.

**Writing – original draft:** Dongqiang Wang, Meng Li, Jigang Yin, Guan Zhu.

**Writing – review & editing:** Guan Zhu.

